# mTOR suppresses macroautophagy during postnatal development of the striatum

**DOI:** 10.1101/536680

**Authors:** Ori J. Lieberman, Irena Pigulevskiy, Michael R Post, David Sulzer, Emanuela Santini

## Abstract

Macroautophagy (hereafter referred to as autophagy) plays a critical role in neuronal function related to development and degeneration. Here, we investigated whether autophagy is developmentally regulated in the striatum, a brain region implicated in neurodevelopmental disease. We demonstrate that autophagic flux is suppressed during striatal postnatal development, reaching adult levels around postnatal day 28 (P28). We also find that mTOR signaling, a key regulator of autophagy, increases during the same developmental period. We further show that mTOR signaling is responsible for suppressing autophagy, via regulation of Beclin-1 and VPS34 activity. These results demonstrate that neurons coopt metabolic signaling cascades to developmentally regulate autophagy and establish mTOR as a central node in the regulation of neuronal autophagy.

## Introduction

Originally characterized in studies of brewer’s yeast, macroautophagy (hereafter referred to as autophagy) is a degradative process for long-lived proteins and damaged organelles (Ohsumi, 2014). In neurons, autophagy is considered to play both protective and pathogenic roles, as it contributes to proteostasis and cell survival (Komatsu et al., 2006; Hara et al., 2006; Yamamoto et al., 2006; Yamamoto and Simonsen, 2011), while some neurons are thought to undergo autophagic cell death in neurodegenerative diseases (Yang et al., 2011; Yamamoto and Yue, 2014; González-Polo et al., 2005; Chakrabarti et al., 2009; Wang et al., 2006). Recently, autophagy has been recognized as an important cellular process during neuronal development (Ebrahimi-Fakhari et al., 2016a; Lieberman et al., 2018b; Dragich et al., 2016; Tang et al., 2014; Kim et al., 2017). Autophagic dysfunction is observed in neurodevelopmental disorders (Poultney et al., 2013; Hor and Tang, 2018; Byrne et al., 2016; Lee et al., 2013), and mouse models with reduced autophagy display phenotypes implicated in autism spectrum disorders (ASD) (Tang et al., 2014; Kim et al., 2017; Yan et al., 2018). Yet, little is known about the developmental regulation of neuronal autophagy.

The striatum is the main input nucleus of the basal ganglia, a brain circuit controlling action selection and reward processing, and is implicated in the pathophysiology of multiple neurodevelopmental diseases (Gerfen and Surmeier, 2011; Fuccillo, 2016). Although the principal neurons of the striatum, the spiny projection neurons (SPNs), migrate to the striatum during the embryonic period (Song and Harlan, 1994), significant further maturation occurs during the first four postnatal weeks. SPNs receive excitatory inputs from the cortex and thalamus during the second and third postnatal weeks (Tepper et al., 1998). Dopaminergic axons innervate the striatum at birth but their ability to release neurotransmitter increases during the first two weeks (Voorn et al., 1988; Lieberman et al., 2018a). Finally, the intrinsic excitability of SPNs matures from weeks two through four (Tepper et al., 1998). Autophagy has been proposed to contribute to synaptic maturation and plasticity and dopamine release (Hernandez et al., 2012; Tang et al., 2014; Nikoletopoulou et al., 2017). Thus, establishing whether autophagy is differentially regulated during postnatal development of the striatum would provide an insight into its role in neurodevelopment.

Autophagy is a tightly regulated multi-step process (Bento et al., 2016) that, in dividing cells is controlled by energy balance (i.e., nutrient status) and metabolic kinases, including the mammalian target of rapamycin (mTOR). mTOR regulates autophagy via several mechanisms (He and Klionsky, 2009), including by phosphorylating and negatively regulating Unc-51-like autophagy-activating kinase 1 (ULK1) at Ser757 (Kim et al., 2011; Jung et al., 2009). This step prevents ULK1-mediated phosphorylation of Beclin-1 at Ser14, and the subsequent increase of PI3K activity of Vps34 (Russell et al., 2013). These molecular events initiate the formation of preautophagic structures that are subsequently expanded by a molecular cascade resulting in the modification of LC3, one of the mammalian homologs of the yeast Atg8 (Shpilka et al., 2011). Processing of LC3, leads to phagophore expansion and sealing and is used as a biochemical readout of autophagosome formation (Kabeya et al., 2000; Klionsky et al., 2016). The enclosed, mature autophagosome then traffics to the lysosome where the autophagic cargo and cargo adaptors, such as p62, are degraded (Tanida et al., 2005).

Whether similar signaling regulates autophagy in neurons remains controversial. It has been proposed that autophagy may act as a constitutive process for cellular homeostasis, thus circumventing the control of metabolic kinases such as mTOR (Yamamoto and Yue, 2014). The links between nutrient status and autophagy in neurons moreover remain elusive, with reports suggesting a regional and age-specific autophagic response to nutrient deprivation in neurons (Kaushik et al., 2011; Nikoletopoulou et al., 2017) and studies indicating the contrary (Mizushima et al., 2004). Moreover, direct regulation of autophagy by mTOR, independent of nutrient status, has been reported by some (Hernandez et al., 2012; Tang et al., 2014) but not others not (Maday and Holzbaur, 2016; Tsvetkov et al., 2010).

These contrasting results raise the important question of whether autophagic activity is dynamically controlled in brain regions implicated in neurodevelopmental disease, and if so, whether autophagy in neurons is regulated by the same pathways that control nutrient deprivation-induced autophagy in nonneuronal cells.

Using biochemical, pharmacological and histological approaches, we demonstrate that autophagy is dynamically downregulated during postnatal development, following the upregulation of mTOR activity. Our results suggest that autophagy may play temporally-specific roles in brain development.

## Results

### Markers of autophagic activity decrease during postnatal development

To identify changes in autophagic activity during postnatal striatal development, we collected striata from mice at postnatal days 8, 14, 18, 28 and in adults (postnatal day 120; (Tepper et al., 1998; Lieberman et al., 2018b; Peixoto et al., 2016). These postnatal ages represent critical timepoints for striatal development. Briefly, synaptic dopamine release has begun and interneurons have migrated in the striatum at postnatal day 8 (Plotkin et al., 2005; Lieberman et al., 2018b; Ferrari et al., 2012). By postnatal day 14, excitatory inputs from the cortex and thalamus arrive and eye opening has occurred (Tepper et al., 1998), providing higher levels of sensory input. P18 represents the end of synaptogenesis and an age immediately before weaning (Tepper et al., 1998). At age P28, the period of postnatal refinement has ended (Tepper et al., 1998). We compared tissue from these ages to mice in early adulthood at postnatal day 120.

We first measured the levels of DARPP32, a classic SPN marker, and actin as a loading control across postnatal development and found no differences (Figure 1A; (Tanida et al., 2005; Klionsky et al., 2016). We then measured the level of total and processed form of the Atg8 family member, LC3B. We found a significant effect of age on the levels of processed LC3B (LC3B-ii) relative to actin and unprocessed LC3B (LC3B-i) (Figure 1A-C). The level of the autophagic adapter protein, p62, whose steady-state levels are determined by its own autophagic degradation, increased over the postnatal period (Figure 1D). These data suggest that overall autophagic activity decreases during the first four postnatal weeks.

**Figure 1.**
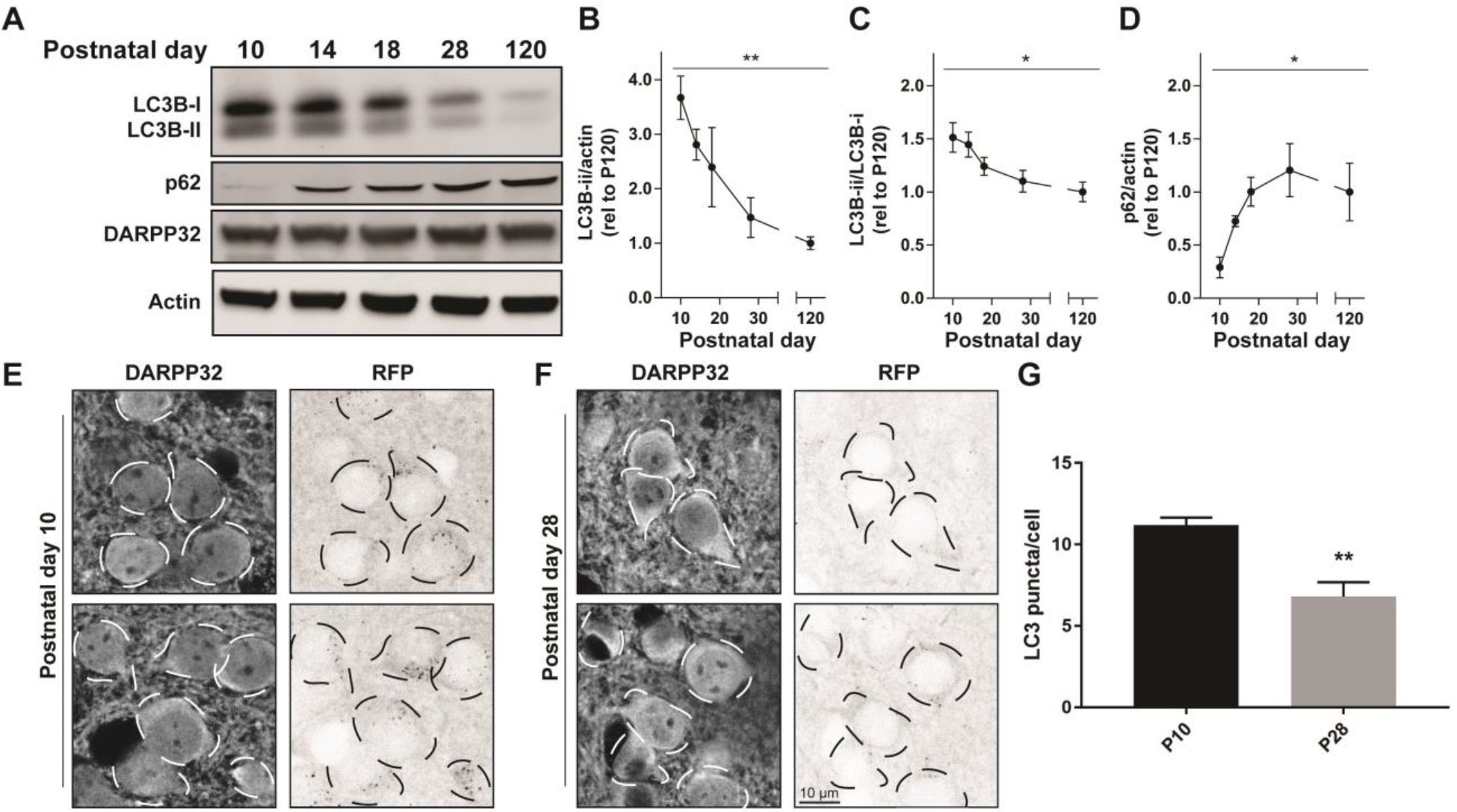
Autophagy decreases during striatal development. **(A)** Representative Western blot images for LC3B, p62, DARPP32, and actin. **(B-D)** Quantification of **(B)** LC3B-ii relative to actin, **(C)** LC3B-ii relative to LC3B-i, **(D)** p62 relative to actin normalized to P120 values. Data analyzed with one-way ANOVA; (B) Age: F_(4,21)_ = 6.526, p=0.0014; (C) Age: F_(4,22)_=3.797, p=0.0171; (D) Age: F_(4,15)_=3.762, p=0.0260. * P<0.05, ** p< 0.01, **** p<0.0001, n=4-6 mice/ age. **(E-F)** Representative images of DARPP32 stained striatal neurons and RFP fluorescence from mice aged P10 and P28. Dashed lines indicate cell body outlines. **(G)** Quantification of number of LC3 puncta/cell. Unpaired, two-tailed t test, t_4_ = 4.392, p=0.0118. N=3 mice/ age.

### LC3+ puncta decrease in striatal spiny projection neurons during postnatal development

Western blot analysis of total striatal lysates includes proteins from all cell types present in the striatum, including neurons, glia, and vascular cells. To define the cell type in which the developmental changes in autophagy occur, we utilized a transgenic mouse ubiquitously expressing LC3 fused to both green (GFP) and red fluorescent proteins (RFP) (Figure S1A; tandem fluorescent-tagged LC3 or tfLC3) (Li et al., 2014). After processing, LC3 transitions from the cytosol to become membrane-bound on the autophagosome (LC3B-ii). Visualizing the distribution of fluorophore-tagged LC3 (LC3 puncta) provides a well-established assay for monitoring autophagic activity within a cell (Klionsky et al., 2016). Furthermore, as the fluorescence of the GFP component is quenched by the low pH of the lysosome, tfLC3 permits analysis of the total number of autophagosomes and autolysosomes (RFP+ puncta) and non-acidified autophagosomes (GFP+RFP+ puncta).

We first used 2-photon microscopy to simultaneously image GFP and RFP signals in acute brain slices tfLC3 mice at age P10 and P28 (Figure S1B-C). At both ages, GFP fluorescence was diffuse in the cytosol and processes of striatal cells and RFP+ puncta were present in the soma. The density of the striatal GFP fluorescence was too strong in the soma to discern individual puncta, we used the number of RFP puncta as a proxy for autophagosomes, with the caveat that we were unable to determine whether these puncta were also fused with lysosomes.

To identify the number of RFP+ puncta in specific cell types, we perfused mice at age P10 and P28 and co-labelled with cell-type specific markers of striatal neurons. Spiny projection neurons can be identified by immunostaining with antibodies against DARPP-32. We observed a significant decrease in RFP+ puncta in DARPP32+ cells between P10 and P28 (Figure 1E-G), indicating that the reduction in autophagic activity we measured using biochemical assays of bulk striatal lysates occurs specifically in SPNs.

### Autophagosome biosynthesis is suppressed during postnatal development

Changes in the level of LC3B-ii, or the number of LC3 puncta, can arise from increases in autophagosome biosynthesis or decreases in the efficiency of autophagosome maturation and lysosomal degradation. To dissect the changes in autophagy during the postnatal period, we developed an *ex vivo* system to test the effects of drugs that do not cross the blood-brain barrier (Figure 2A). We generated acute brain slices from P10 or P28 mice and removed non-striatal tissue. We found that neither the slice procedure nor the incubation affected the age-dependent reduction in autophagy markers observed *in vivo* (Figure 2A-B compared to Figure 1). This confirms that the mechanisms underlying changes in autophagic markers during postnatal development can be defined using this *ex vivo* system.

**Figure 2.**
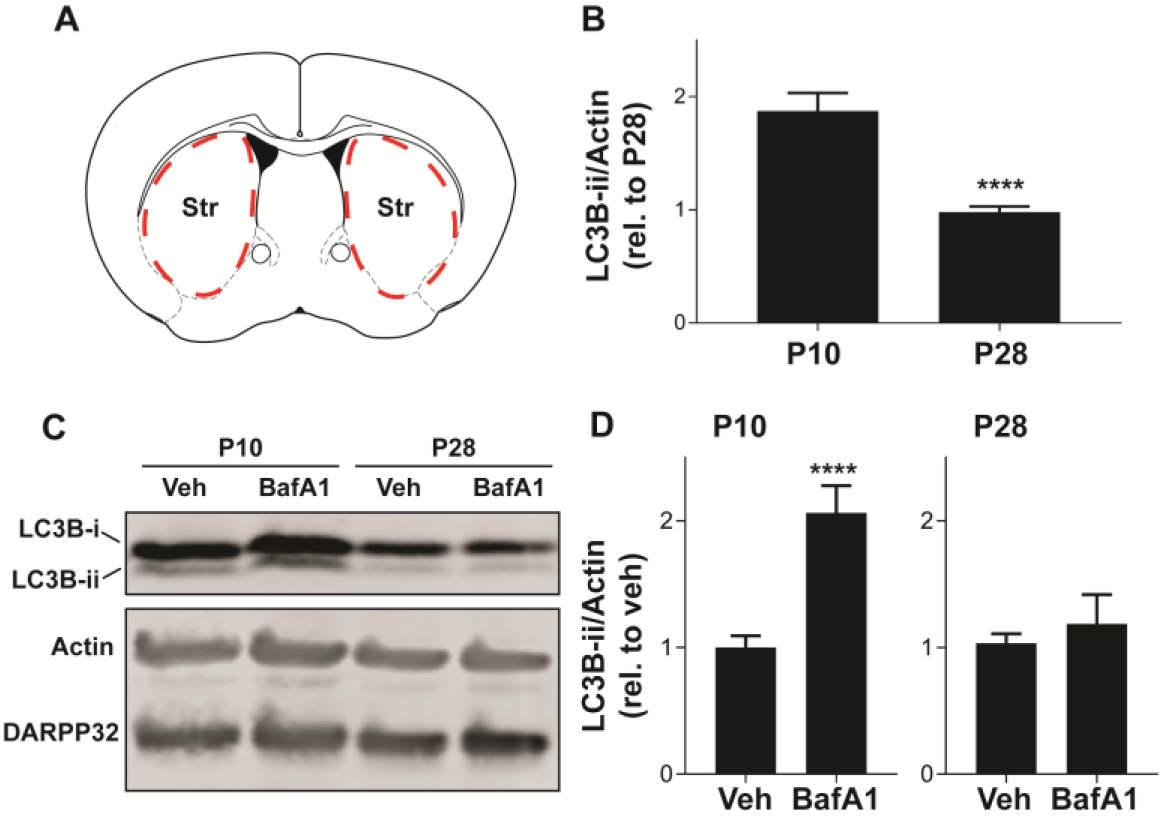
Striatal autophagosome biosynthesis decreases from P10-P28. **(A)** Schematic of coronal brain section showing dissection boundaries for ***ex vivo*** experiments. **(B)** Quantification of LC3B-ii relative to actin for every vehicle-only slice showed in Figures 2, 4, and 5. Unpaired, two-tailed t test, t52 = 5.824, ****p<0.0001. **(C)** Representative Western blot images of actin, DARPP32 and LC3B in slices obtained from P10 or P28 mice, incubated with BafA1 (100 nM, 3 hours) or vehicle (Veh; DMSO, 0.1%). **(D)** LC3B-ii relative to actin, normalized to vehicle condition at each age. P10: Unpaired, two-tailed t test, t25 = 5.113, ****p<0.0001; P28: Unpaired, two-tailed t test, t10 = 0.6228, p=0.5473. P10: Veh: n= 16 slices, BafA1 n = 11 slices from 46 mice. P28: Veh: n=6 slices, BafA1 n=6 slices from 3 mice.

We incubated slices from mice at age P10 and P28 with bafilomycin A1 BafA1, 100 nM), a specific inhibitor of the vacuolar proton pump, or vehicle (DMSO, 0.1%) for three hours, an incubation time similar to that used in cultured cells (Klionsky et al., 2016), to prevent lysosomal acidification and block autophagosome-lysosome fusion. As treatment with BafA1 prevents LC3B-ii degradation, changes in LC3B-ii levels following BafA1 treatment are interpreted as the rate of autophagosome biosynthesis. BafA1 treatment increased LC3B-ii in slices from P10 mice, but had no significant effect in slices from P28 mice (Figure 2C-D). BafA1 treatment had no effect on DARPP32 levels at either age (Figure 2C). Slices from either age treated with BafA1 for one hour showed no change in LC3B-ii levels (data not shown). This suggests that the higher baseline level of LC3B-ii at P10 arises from increased autophagosome biosynthesis.

### mTOR signaling is upregulated during the postnatal development

mTOR signaling is a key negative regulator of autophagic activity. We therefore hypothesized that mTOR activity in the striatum increases during postnatal development and suppresses autophagy.

mTOR kinase activity can be monitored by measuring the state of phosphorylation of its downstream targets. The level of phosphorylation at serine 757 of ULK1, which is phosphorylated by mTOR and inhibits ULK1 kinase activity (Kim et al., 2011), increased during the postnatal period in the striatum (Figure 3A-C). We then confirmed that ULK1 kinase activity is inhibited by monitoring the phosphorylation state of Beclin-1, a ULK1 target. Phosphorylation of Beclin-1 at serine 14 decreased during postnatal development (Figure 3A-C). mTOR activity also leads to the phosphorylation of the ribosomal protein S6 (rpS6) on serine 240 and serine 244 (Magnuson et al., 2012). We observed a sharp increase in rpS6 Ser240/244 at P18 before decreasing into adulthood (Figure 3B and 3D). Overall these data indicate an increase in mTOR activity during striatal postnatal development. Interestingly, we did not observe a significant effect of age on phosphorylation of ERK1/2 (Figure 3B and 3D) suggesting that increased mTOR activity during the postnatal period is not associated with a global change in molecular signaling.

**Figure 3.**
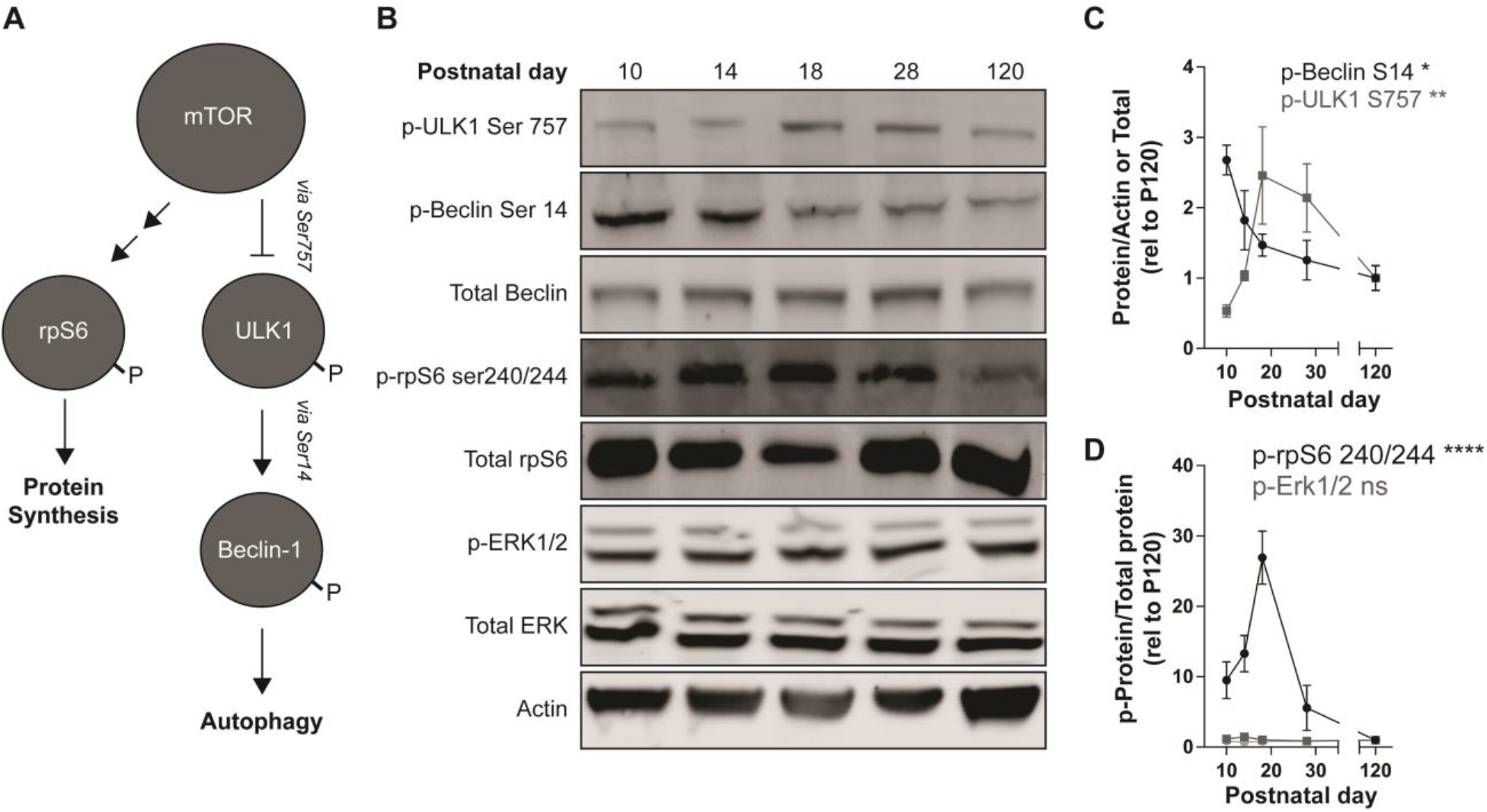
mTOR signaling increases during striatal development. **(A)** Schematic representation of mTOR targets. mTOR inhibits ULK1 activity by phosphorylating Ser757. P-ULK1 activates Vps34 activity (not shown) by phosphorylating Beclin-1 on Ser14. mTOR promotes protein synthesis by phosphorylating rpS6 on Ser240/244. **(B)** Representative Western blot images quantified in C-D. **(C)** Quantification of pULK1 S757 relative to actin (grey squares) and p-Beclin S14 relative to total Beclin-1 (black dots). Data analyzed with one-way ANOVA; p-ULK1 S757/ actin: Age: F_(4,13)_ = 6.093, p=0.0055; p-Beclin-1S14/ total Beclin-1: Age: F_(4,14)_ = 4.945, p=0.0107. **(D)** Quantification of p-rpS6 S240/244 and p-Erk1/2 relative to total rpS6 and total Erk1/2, respectively. Data analyzed with one-way ANOVA; p-rpS6 240/244: Age: F_(4,20)_ = 12.69, p<0.0001. p-Erk1: Age: F_(4,17)_x = 2.625, p=0.0712. p-Erk2: Age: F_(4,17)_ = 0.3561, p=0.8362. N=3-6 mice/ age. * p<0.05, ** p<0.01, **** p<0.0001.

### Inhibition of Vps34 reduces LC3B-ii levels during early postnatal development

mTOR negatively regulates autophagosome biosynthesis by inhibiting ULK1 activity (Jung et al., 2009; Kim et al., 2011). When ULK1 is active, it promotes autophagosome formation by phosphorylating Beclin-1, which increases the PI3K activity of its partner, Vps34 (Russell et al., 2013). To address whether elevated autophagy at P10 was a result of enhanced Vps34 activity, we incubated acute striatal slices with the Vps34 inhibitor, SAR405 (1 μM) or vehicle (DMSO, 0.1%) for three hours (Ronan et al., 2014). SAR405 significantly reduced the level of LC3B-ii in slices from mice at age P10 but had no effect on slices from P28 mice (Figure 4A-B). This demonstrates that elevated autophagic activity in the striatum of early postnatal mice is Vps34-dependent. The lack of effect of SAR405 on autophagic activity at P28 indicates that reduced Vps34 activity, possibly via increased mTOR signaling (Figure 3), is responsible for the lower levels of autophagosome biosynthesis at P28.

**Figure 4.**
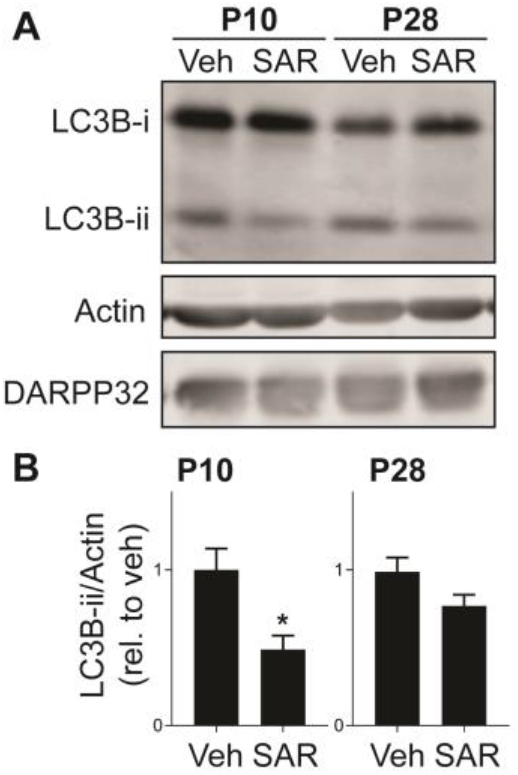
Vps34 activity is required for maintaining LC3B-ii levels at P10. **(A)** Representative Western blot images for actin, DARPP32 and LC3B-i and -ii in striatal slices obtained from P10 or P28 mice, treated with SAR405 (1 μM) or vehicle (Veh; DMSO, 0.1%). **(B)** Quantification of LC3B-ii relative to actin, normalized to vehicle condition at each age. P10: unpaired, two-tailed t test, t_11_ = 2.985, p=0.0124; P28: Unpaired, two-tailed t test, t_24_ = 1.807, p=0.0922. P10: Veh: n=6 slices, SAR405 n=7 slices from 3-4 mice/ age. P28: Veh: n=9 slices, Baf n=7 slices from 3-4 mice/ age.* p<0.05.

### mTOR inhibition increases LC3B processing during the late postnatal period

Having shown that Vps34 drives elevated autophagy at P10, we explored whether enhanced mTOR activity inhibits autophagy at P28. mTOR activity can be pharmacologically inhibited by direct active site inhibitors, such as Torin-1 (Thoreen et al., 2009). We incubated striatal slices from mice at P28 with Torin-1 (5 μM) or vehicle (DMSO, 0.1%) for three hours and measured LC3B-ii levels. Torin-1 treatment increased LC3B-ii levels and the phosphorylation of Beclin-1 at the ULK1 site (Ser14), suggesting that mTOR inhibition activates autophagy in a ULK1/Beclin-1 dependent manner (Figure 5A-C).

**Figure 5.**
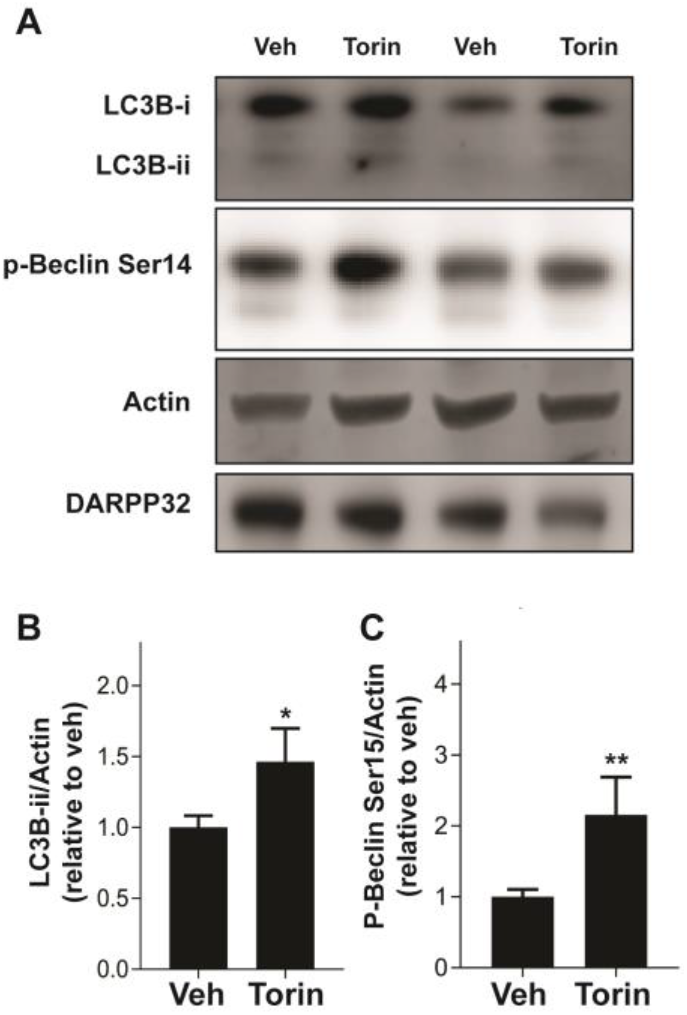
mTOR inhibition increases p-Beclin-1 and LC3B-ii levels at P28. **(A)** Representative Western blot images for actin, DARPP32, p-Beclin-1 Ser14 and LC3B-i and -ii in striatal slices from P28 mice, treated with Torin-1 (5 μM) or vehicle (Veh; DMSO, 0.1%). **(B-C)** Quantification of **(B)** LC3B-ii and **(C)** p-Beclin S14 relative to actin, normalized to vehicle. Data analyzed with unpaired, two-tailed t test; **(B)** LC3B-ii/ actin, t_34_ = 2.312, p=0.0270. Veh: n=24 slices, Baf n=12 slices from 5-7 mice.* p<0.05. **(C)** p-Beclin S14/ actin, t_22_ = 3.337, p=0.0030. ** P<0.01, Veh: n=18 slices, Torin-1 n=6 slices from 3-5 mice.

## Discussion

Neuronal autophagy has been proposed to play a key role in neurodevelopment and autophagic dysfunction may lead to neurodevelopmental disorders (Lieberman et al., 2018b; Tang et al., 2014; Yan et al., 2018; Kim et al., 2017; Dragich et al., 2016). In this study we address whether autophagic activity is developmentally controlled in the principal neurons of the striatum, a brain region implicated in neurodevelopmental disorders (Fuccillo, 2016). We found that autophagy is dynamically regulated in SPNs during postnatal development, reaching adult levels around P28. These findings provide mechanistic insight into the regulation of autophagy during striatal postnatal development and establish the basis for a further evaluation of its role in physiological and pathological conditions.

By analyzing key endogenous biochemical markers (such as LC3B-ii, p62, p-Ser757-ULK1 and p-Ser14-Beclin-1), we discovered that autophagy decreases progressively during the postnatal development of SPNs. The autophagy markers were measured in bulk lysates containing neurons, glia and endothelial cells (among others). To identify the cell types that feature changes during postnatal development, we utilized tfLC3 mice (Li et al., 2014). Given the challenges in assaying endogenous autophagic proteins using immunofluorescence, tfLC3 mice provide a unique resource, as overexpressed LC3 is fused to fluorescent proteins that permit morphological characterization of autophagic structure. While the high levels of expression prevented the identification of individual autophagic puncta in both live imaging (Figure S1) and fixed tissue (not shown), the level of autophagolysosomes represented by RFP-LC3+ puncta was higher in SPN somata at P10 than at P28, in agreement with our biochemical analysis (Figure 1).

Autophagic flux is determined by the kinetics of autophagosome biosynthesis, autophagosome maturation and autophagic cargo degradation by lysosomal proteases. Steady-state measurement of LC3B-ii levels provide a useful proxy for the measurement of autophagic activity. An elevated level of LC3B-ii could indicate either increased or decreased autophagic flux, as both increased autophagosome biosynthesis *and* decreased lysosomal degradation of autophagic proteins can increase LC3B-ii (Kabeya et al., 2000; Klionsky et al., 2016). To functionally measure autophagic flux in striatal tissue, we developed an *ex vivo* acute brain slice system and found that steady state increases in LC3B-ii levels at P10 were further elevated by acute blockade of lysosomal degradation with BafA1 compared to LC3B-ii levels at P28. This suggests that increased autophagosome biosynthesis is responsible for the higher steady-state LC3B-ii levels, as inhibition of lysosomal degradation would not have an effect on LC3B-ii levels if lysosomal degradation were already compromised. These results indicate that the postnatal suppression of autophagy results from changes in autophagosome biosynthesis. Determining whether autophagosome formation is critical for the degradation of specific cargo during the early postnatal period or whether this represents a key switch in modes of membrane trafficking will be important to understanding the role of autophagy in neurodevelopment.

In line with changes in autophagosome biosynthesis during striatal maturation, we found that mTOR-dependent signaling increases over the course of postnatal development, thereby suppressing autophagosome synthesis via inhibition of ULK1 and Beclin-1 activity. We also determined that the phosphorylation of mTOR targets increases during the third postnatal week. These include downstream targets involved in protein synthesis, such as rpS6, and autophagy, such as ULK1. mTOR specifically phosphorylates ULK1 at Ser757 and inhibits its kinase activity (Kim et al., 2011). Consistently, we also observed a decrease in the level of phosphorylation on the ULK1 site in Beclin-1, a key component of the class III-PI3K complex required for autophagosome biosynthesis (Russell et al., 2013; Itakura et al., 2008). These results provide a functional readout for increased mTOR activity on autophagy-specific downstream targets and demonstrate a direct link between mTOR signaling and a regulation of autophagy activity observed during the postnatal development. Inhibition of mTOR activity at P28, when mTOR activity is elevated, induces autophagy and increases the phosphorylation of Beclin-1 at Ser14 and LC3B-ii levels. Further work will also focus on whether additional kinases which integrate metabolic status in the periphery, such as AMPK (He and Klionsky, 2009), control developmental changes in neuronal autophagy.

These findings may help to resolve conflicting results present in the literature. Several groups have found that mTOR inhibition fails to increase autophagic activity in primary neuronal culture (Maday and Holzbaur, 2016; Tsvetkov et al., 2010) or *in vivo* (Fox et al., 2010). In contrast, others report that pharmacological (Hernandez et al., 2012; Tang et al., 2014) or genetic inhibition of mTOR drives autophagy (Yan et al., 2018) as well as hyperactivation of mTOR signaling inhibits autophagy in the CNS (McMahon et al., 2012; Tang et al., 2014; Ebrahimi-Fakhari et al., 2016b). This discrepancy may arise from model system used (i.e., cell culture vs *in vivo*), developmental stage, or treatment paradigm. Further complicating the link between mTOR and autophagy is that standard activators of autophagy such as starvation *in vivo* and nutrient or serum deprivation *ex vivo* may not have effects in the brain (Maday and Holzbaur, 2016; Young et al., 2009; Mizushima et al., 2004; Nikoletopoulou et al., 2017). Our findings suggest that mTOR signaling is important for the regulation of autophagy during critical developmental windows in neurons in a manner unrelated to nutrient status. mTOR signaling is under the control of several neural-specific signals such as patterned neuronal activity and specific neurotransmitters or neuromodulators (Sutton and Caron, 2015; Santini et al., 2009; Auerbach et al., 2011; Yin et al., 2006; Bockaert and Marin, 2015), suggesting possible mechanisms through which mTOR activity is regulated within developing neural circuits.

In summary, we find that the postnatal development of SPNs is correlated with a change in autophagy and this is determined by mTOR signaling status. Intriguingly, this signaling cascade is also critical for nutrient-deprivation induced autophagy in dividing cells (He and Klionsky, 2009), suggesting that the same signaling cascade may be triggered by different cues depending on the tissue type. There is a broad range of significant changes in neurotransmitter content, neuronal firing patterns and synaptic plasticity that occurs during this period, providing a plethora of candidates that could dynamically regulate mTOR signaling and autophagy during the course of postnatal development. Autophagy is also known to regulate surface levels of neurotransmitter receptors, axon pathfinding, synaptic maturation and plasticity, suggesting that autophagy acts dynamically during early postnatal developmental to entrain adult-like neuronal activity that subsequently suppresses autophagy by activating mTOR signaling (Rowland et al., 2006; Shehata et al., 2012; Dragich et al., 2016; Nikoletopoulou et al., 2017; Tang et al., 2014; Hernandez et al., 2012). Such a feedback process may provide a temporal and mechanistic framework for future studies that address the role of autophagy, and its regulation by mTOR, in neuronal development.

## Methods

### Animals

Breeder pairs of C57/BL6J were obtained from Jackson Laboratories (Bar Harbor, ME). Mice were checked every day or every other day for pregnancy and new litters. TfLC3 mice were obtained from Jackson Laboratories (C57BL/6-Tg(CAG-RFP/EGFP/Map1lc3b)1Hill/J; Strain No. 027139). Breeder pairs were housed on a 12-hour light/dark cycle with water and food available *ad libitum*. Offspring were weaned between postnatal day 18 and 20 and split into same sex groups of 2-5. All experimental procedures were approved by the Columbia University Institutional Animal Care and Use Committee. Mice of both sexes were utilized for experiments and data were combined as no effect of gender was observed.

### *In vivo* sample preparation

Striatal sample preparation was performed as previously described (Santini et al., 2007). Mice at specified ages were rapidly decapitated and their head was briefly placed in liquid nitrogen. The brain was subsequently removed and a single striatum was dissected and flash frozen in liquid nitrogen. Samples were stored at −80 °C until the full cohort was collected. Samples were then homogenized in 1% SDS by a brief sonication. Protein content was determined by the BCA assay (Thermo Fisher). No significant effect of age was found on total protein (data not shown). Samples were then boiled in sample buffer and frozen until western blot analysis.

### Acute brain slice

Acute brain slices were prepared essentially as described (Lieberman et al., 2018a). Briefly, mice underwent cervical dislocation and the brain was removed and placed in ice-cold high sucrose cutting solution (in mM): 10 NaCl, 2.5 KCl, 25 NaHCO_3_, 0.5 CaCl_2_, 7 MgCl_2_, 1.25 NaH_2_PO_4_, 180 sucrose, 10 glucose bubbled with 95% O_2_/5% CO_2_ to pH 7.4. Brains were mounted on a VT1200 vibratome (Leica Biosystems) and coronal sections (250 μm) including the striatum were collected. Slices were then transferred to a basin contained ice-cold cutting solution and the striatum was manually dissected. Slices were moved to scintillation vials containing 7 mL of ACSF (in mM): 125 NaCl, 2.5 KCl, 25 NaHCO_3_, 2 CaCl_2_, 1 MgCl_2_, 1.25 NaH_2_PO_4_, and 10 glucose bubbled with 95% O_2_/5% CO_2_ to pH 7.4 at 34 °C. Slices from two mice at the same age were combined in individual experiments and split into 4 conditions. Slices were allowed to rest for one hour followed by addition of vehicle or drug. Following incubation with drug at 34 °C for the specified time, slices were removed and flash frozen in liquid nitrogen. After slices were collected from a complete experiment (i.e. slices from 4-6 mice per age, at P10 and P28), slices were homogenized by sonication in 1% SDS and prepared as described above for Western blot analysis.

### Western blot

Equivalent amount of protein per sample (5-25 μg/well) were loaded into 10% or 12% polyacrylamide gels as described (Santini et al., 2007). Protein was transferred from the gel to an Immobilon FL PVDF membrane (pore size 0.2 μm). Blots were blocked in TBS with 0.1% Tween-20 (TBST) and 5% bovine serum albumin (BSA) for one hour at room temperature. Blots were then incubated with primary antibody (see Table 1 for detailed information regarding antibodies) diluted in TBST with 5% BSA as specified (see table 1 for dilution and antibody sources) overnight at 4 °C. Blots were then washed with TBST and incubated in secondary antibody for one hour at room temperature. Blots were developed using either the Odyssey imaging system (LICOR) or an enhanced chemiluminescence (ECL) system (Amersham) and imaged using an Azure Biosystems C600 system. Western blots developed using the Odyssey system were analyzed in Image Studio Lite (LICOR). Western blots developed with the ECL system were analyzed using standard routines in ImageJ. All samples were probed for beta-actin and DARPP32 as loading and dissection controls, respectively.

**Table 1.** Antibodies

| Antibody | Company<br>(Catalog No.) | Dilution | Detection Method | Notes |
| --- | --- | --- | --- | --- |
| Rabbit anti LC3B | Novus Biologicals<br>(NB600-1384) | 1:1000 | Odyssey | 12% SDS-PAGE<br>gel required |
| Rabbit anti p62 | MBL (PM045) | 1:1000 | Odyssey | Atg7KO<br>validated (data<br>not shown) |
| Rabbit anti<br>Darpp32 | Cell Signaling<br>Technology<br>(2306) | 1:2500-5000 | Odyssey |  |
| Mouse anti β-<br>Actin | Novus Biologicals<br>(NB600-501) | 1:5000-10000 | Odyssey | |
| Rabbit anti p-ULK1 (Ser757) | Cell Signaling<br>Technology<br>(6888) | 1:500 | Odyssey |  |
| Rabbit anti Beclin | Cell Signaling<br>Technology<br>(3495) | 1:1000 | ECL |  |
| Rabbit anti p-Beclin-1 (Ser14) | **Cell Signaling Technology (84966)** | **1:1000** | **ECL** |  |
| Rabbit anti S6 ribosomal protein | **Cell Signaling Technology (2217)** | **1:1000-2000** | **ECL** |  |
| Rabbit anti p-S6 ribosomal protein (Ser240/244) | **Cell Signaling Technology (2215)** | **1:1000-2000** | **ECL** |  |
| Rabbit anti p44/42 MAPK (Erk1/2) | **Cell Signaling Technology (4695)** | **1:1000** | **Odyssey** |  |
| Rabbit anti p-p44/42 MAPK (Erk1/2; Thr202/Tyr204) | **Cell Signaling Technology (4370)** | **1:1000** | **Odyssey** |  |

### Drugs and chemicals

BafA1 (0.1 μM) and Torin-1 (5 μM) were purchased from Tocris. SAR 405 (1 μM) was purchased from Cayman Chemicals. All drugs were dissolved in DMSO and slices were incubated in ACSF containing the drugs or equivalent volume of vehicles for three hours at 34°C. DMSO did not exceed a final concentration of 0.1%. All other chemicals were purchased from Fisher Scientific.

### 2p-microscopy

Two-photon images were acquired on a Prairie Ultima microscope system (Middleton, WI) using PrairieView 4.3 software. Acute brain slices were transferred into a chamber and perfused with oxygenated ACSF at room temperature. Samples were excited with a Coherent (Santa Clara, CA) Chameleon Ultra two-photon laser at 980 nm, and images were simultaneously collected through two photomultiplier tube channels with corresponding 585-630 nm and 490-560 nm emission windows. The objective used was a 60X, 0.9 NA water immersion lens, and images were 1024×1024 pixels in size.

### Immunohistochemistry

Mice were deeply anesthestized and transcardially perfused with 0.9% Nacl followed by 4% paraformaldehyde (PFA) in 0.1M phosphate buffer (PB). Brains were removed and post-fixed overnight in 4% PFA in 0.1M PB. Brains were then washed three times in 1X phosphate buffered saline and cut into 40 μm sections using a VT1200 vibratome (Leica Biosystems) and stored in cryoprotectant (0.1M PB, 30% glycerol, 30% ethylene glycol) at −20 °C. For immunofluorescence analysis, sections were washed in TBS three times and then blocked and permeabilized for one hour at room temperature with 10% normal donkey serum (Jackson Immunoresearch) and 0.1% Triton-X in TBS. Sections were then incubated overnight at 4 °C with primary antibodies in 2% normal donkey serum, 0.1% Triton-X in TBS. Primary antibodies included: Rabbit anti-DARPP32 (Cell Signaling), and Chicken anti Green Fluorescent Protein (Abcam). Secondary antibodies with the appropriate conjugated fluorphores were purchased from Invitrogen. The endogenous fluorescence of RFP was imaged. Sections were then washed in TBS and mounted. Images were obtained using a Leica SP5 confocal system with argon, DPSS He/Ne lasers. Images were obtained with a 63X oil immersion objective with a 2x digital zoom at 2048×2048 resolution (~120 nm resolution).

All images were taken with the same laser intensity and detector settings with non-saturating pixel intensities.

Image analysis was conducted in ImageJ. Cell bodies were segmented using DARPP32 staining. RFP+ puncta within each segment was counted manually. 10-20 cells were counted per section, 2-4 sections were counted per animal. Average number of puncta per cell was determined from n=3 animals per age. All images were collected and all analyses were conducted blind to condition.

### Statistics

All analysis was conducted blind to condition. For statistical analysis between two groups, unpaired, two-tailed t tests were used. For analysis between 3 or more groups, one-way ANOVA was used. Normality was not formally tested. Sample size was not based on a formal power analysis but was based on past work from our groups and similar experiments from the literature. Statistical analysis was conducted in GraphPad Prism 7 (La Jolla, Ca). All bar graphs show the mean ± SEM.

## Acknowledgements

We thank Ai Yamamoto for insightful discussions about the manuscript; Anders Borgkvist for advice on experimental procedures, mentorship and critical discussions; Dritan Agalliu for generously sharing the LICOR Odyssey system; and Gilberto Fisone for critical reading of the manuscript.

OJL was supported by NIH T32 GM007367 and NIH F30 MH114390, MRP by NIH 5T32MH020004, DS by NIH R01 DA007418 and the Simons and JPB Foundations, ES by NIH R00 NS087112, Swedish Research Council (2016-02758), The Knut and Alice Wallenberg Academy Fellowship, The Olle Engkvist Byggmester Foundation, The Karolinska Institute Strategic Research Program in Neuroscience (StratNeuro) and The Karolinska Institute Internal Funds.

## Author Contributions

Conception: OJL and ES; Methodology: OJL, MRP, and ES; Investigation: OJL, IP, MRP, ES; Writing – Original Draft: OJL; Writing – Review and Editing: OJL, IP, MRP, DS, ES; Visualization: OJL; Supervision: ES; Funding Acquisition: DS and ES.

