## Supplementary material for "mTOR suppresses macroautophagy during postnatal development of the striatum": Figure S1

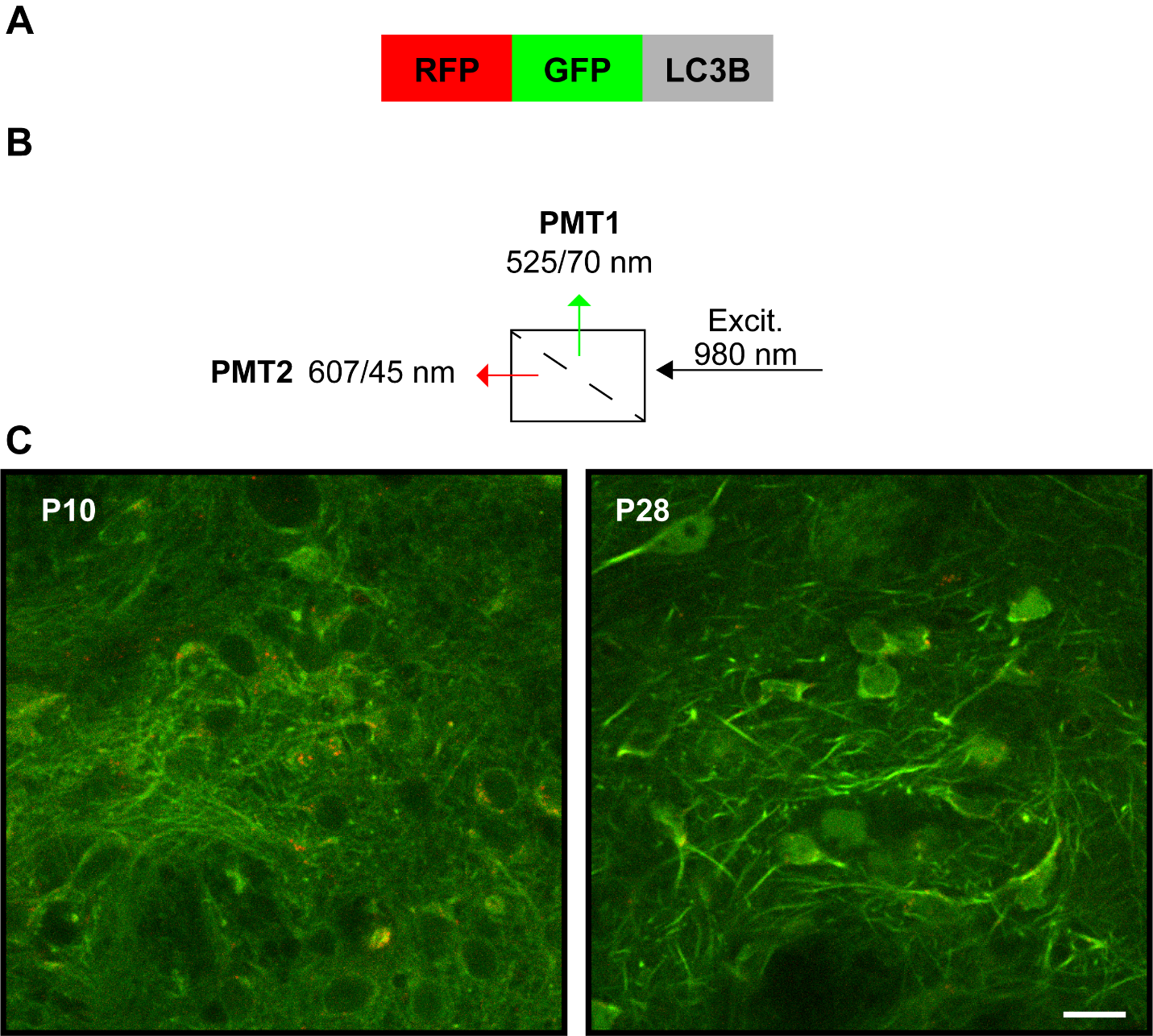

**Figure S1. Live 2-photon imaging of tfLC3 in striatal slice. (A)** Schematic representation of tfLC3 reporter. LC3B fused to RFP and GFP allows detection of cytosolic and membrane bound LC3 by fluorescence microscopy. GFP fluorescence is quenched in acidic environments, permitting analysis of autophagosome and autolysosome numbers. **(B)** Schematic of imaging setup. **(C)** Representative images of tfLC3 mice shows diffuse GFP imaging and punctate RFP patterns at both ages. Scale Bar 25 µm.
